# A common allele in FGF21 associated with preference for sugar consumption lowers body fat in the lower body and increases blood pressure

**DOI:** 10.1101/214700

**Authors:** Timothy M. Frayling, Robin N. Beaumont, Samuel E. Jones, Hanieh Yaghootkar, Marcus A. Tuke, Katherine S. Ruth, Francesco Casanova, Ben West, Jonathan Locke, Seth Sharp, Yingjie Ji, William Thompson, Jamie Harrison, Cecilia M. Lindgren, Niels Grarup, Anna Murray, Rachel M. Freathy, Michael N. Weedon, Jessica Tyrrell, Andrew R. Wood

## Abstract

Fibroblast Growth Factor 21 (FGF21) is a hormone that induces weight loss in model organisms. These findings have led to trials in humans of FGF21 analogues with some showing weight loss and lipid lowering effects. Recent genetic studies have shown that a common allele in the *FGF21* gene alters the balance of macronutrients consumed but there was little evidence of an effect on metabolic traits. We studied a common *FGF21* allele (A:rs838133) in 451,099 people from the UK Biobank study. We replicated the association between the A allele and higher percentage carbohydrate intake. We then showed that this allele is more strongly associated with body fat distribution, with less fat in the lower body, and higher blood pressure, than it is with BMI, where there is only nominal evidence of an effect. These human phenotypes of naturally occurring variation in the *FGF21* gene will inform decisions about FGF21’s therapeutic potential.

## Introduction

FGF21 is a hormone secreted primarily by the liver whose multiple functions include signalling to the paraventricular nucleus of the hypothalamus to suppress sugar and alcohol intake[1, 2], stimulating insulin-independent glucose uptake by adipocytes[3] and acting as an insulin sensitizer [4]. These features and several other lines of evidence have prompted the development of FGF21 based therapies as potential treatments for obesity and type 2 diabetes, with consistent effects on triglyceride lowering, some effects on weight loss but little effect on glucose tolerance [5, 6]. An early trial showed lipid lowering effects in people with type 2 diabetes and obesity but there was only suggestive evidence for effects on weight and glucose tolerance [7]. A recent study suggested that FGF21 analogues may alter blood pressure in humans [8], although changes in blood pressure were not observed in a previous trial [9]. Pre-clinical evidence of FGF21’s potential role in metabolism includes resistance to diet induced obesity in mice overexpressing FGF21[3] and improved glucose tolerance in obese mice through administration of recombinant FGF21[3]. Subsequent studies have confirmed these findings in mice[10] and shown similar effects in non human primates, including improvement of glucose tolerance and slight weight loss in diabetic rhesus monkeys [11], but other studies are less conclusive [5].

Recent studies have shown that FGF21 affects the balance of macronutrients consumed. Studies in mice and non human primates show that genetically and pharmacologically raising FGF21 levels suppresses sugar and alcohol intake[1, 2]. Three human genetic studies have shown that the minor A allele at rs838133 (A/G, Minor Allele Frequency=44.7%), which results in a synonymous change to the first exon of *FGF21*, is associated with higher carbohydrate and lower protein and fat intake, with no effect on total calorie intake[12–14]. Soberg *et al.* showed that the carbohydrate preference was specific to sugary products, and may also increase alcohol intake [13]. These findings are consistent with data from animal studies showing that FGF21 signals to reward centres in the brain [1, 2]. The human genetic studies found no detectable effect on the risk of type 2 diabetes and only nominal evidence for an effect on BMI[13].

Here we aimed to extend the characterisation of the phenotypes associated with the variant in *FGF21* using 451,000 individuals from the UK Biobank. We reasoned that genotype-phenotype associations of *FGF21* would inform developers of FGF21 based therapies as to their potential beneficial and adverse effects. The use of human genetic information is increasingly viewed as an important step to inform drug development[15]. Importantly, the effect size of a human allele is irrelevant to the potential insight into human therapies as long as the associations are statistically robust. We first replicated the association between the minor allele at rs838133 and higher carbohydrate and lower protein and fat intake. We then provide conclusive evidence that the same allele increases alcohol intake, consistent with findings from animal studies. Finally, we show that this allele is associated with stronger effects on body fat distribution, with less fat primarily in the lower body, and higher blood pressure, than its effects on BMI.

## RESULTS

### The minor allele at FGF21 rs838133 is associated with higher sugar and alcohol intake and lower protein and fat intake in UK Biobank individuals

We first investigated the previously described associations between the *FGF21* rs838133 variant and macronutrient intake, coffee and alcohol intake, and smoking. We used data including that derived from a food frequency questionnaire (FFQ) completed by up to 176,994 UK Biobank participants from amongst 451,099 we defined as of European ancestry. In **table 1** we show how each copy of the minor A allele was associated with higher self report estimates of carbohydrate and alcohol intake and lower fat and lower protein intake. These effects imply very strongly that the rs838133 A allele is a loss of *FGF21* function allele because genetic and pharmacological lowering of FGF21 in animal models, including non human primates, has the same effect on carbohydrate and alcohol preferences[1, 2]. The largest effect on macronutrient intake was with carbohydrate intake, where each A allele raised percentage intake by 0.21%. There was no detectable effect on total energy intake. Any effects on coffee consumption and smoking were minimal in comparison. In **Supplementary table 1** we show the summary characteristics of people in the UK Biobank, including those completing the FFQ and those not completing it.

**Table 1.** Associations between the minor A allele at rs838133 and self reported diet and smoking measures in the UK biobank. Macronutrient intake and fizzy drink intake data are based on FFQs completed by up to 176,994 individuals between one and five times. Fizzy drink intake includes calorie free drinks. Other data are based on questionnaires completed at baseline collection. Effect sizes are standard deviations per A allele. Beta raw refers to effect size based on untransformed variable.

| *Quantitative diet outcome* | *N* | *Beta raw (units)* | *Beta (SD)* | *SE* | *P* |
| --- | --- | --- | --- | --- | --- |
| Alcohol units | 341878 | 0.015 (units) | 0.0147 | 0.0022 | $4 \times 10^{-11}$ |
| Total energy | 176994 | -9.623 (KJ) | -0.0036 | 0.0033 | 0.28 |
| % protein | 176989 | -0.108 (%) | -0.0291 | 0.0034 | $3 \times 10^{-17}$ |
| % carbs | 176989 | 0.206 (%) | 0.0244 | 0.0035 | $2 \times 10^{-12}$ |
| % fat | 176989 | -0.196 (%) | -0.0281 | 0.0347 | $4 \times 10^{-16}$ |
| Fizzy drink consumption | 176994 | -0.001 | -0.0028 | 0.0035 | 0.41 |
| Cups of coffee per day* | 299908 | -0.014 | -0.0076 | 0.0027 | 0.005 |
| *Binary outcomes* | *N yes (no)* |  | *OR (95%I)* | *95% CIs* | *P* |
| Current smoker (yes or no) | 35,946 (204,252) | N/A | 0.997 | (0.980, 1.013) | 0.68 |
| Ever smoker (yes or no) | 170,388 (204,252) | N/A | 0.991 | (0.981, 1.000) | 0.06 |
| Coffee (yes or no) | 299,908 (79,235) | N/A | 0.987 | (0.976, 0.998) | 0.025 |

\*within coffee drinkers, (>10 cups per day collapsed into one group), OR=odds ratio, CIs=confidence intervals, N/A=not applicable

### The minor allele at FGF21 rs838133 is associated with lower body fat percentage and altered body fat distribution, with disproportionately less fat in the lower body

Previous studies reported only nominal evidence for associations between rs838133 and anthropometric measures. Using data from 451,099 UK Biobank participants, we showed that the minor rs838133 allele was associated with lower body fat percentage and altered body fat distribution. The A allele was associated with lower hip circumference and higher waist hip ratio (WHR) but there was only nominal evidence of an effect on BMI. The largest effect was in the lower body, where each A allele lowered hip circumference by approximately 1.0 mm. We also observed that the minor allele was associated with shorter stature (also by ~1 mm per allele), but this effect on reduced growth did not account for the smaller hip circumference (**Table 2**). Each of these associations with anthropometric traits was consistent with previously published GWAS data from the GIANT consortium (**Table 2**). Unlike many other variants altering WHR the effects were very similar in men and women (**Supplementary table 2**).

**Table 2.** Associations between the minor allele at rs838133 and anthropometric and metabolic traits in UK Biobank and published GWAS data where available. UK Biobank data are based on 451,000 individuals of European ancestry corrected for relatedness. Effect sizes are in SDs after inverse normalisation or odds ratios for disease traits.

|  | UK Biobank |  |  |  |  | Published GWAS |  |  |  | Meta analysis |  |  |  |
| --- | --- | --- | --- | --- | --- | --- | --- | --- | --- | --- | --- | --- | --- |
| Anthropo-<br>metric trait | N | Beta raw | Beta SD<br>or OR | SE or<br>95% CI | P | N | Beta SD or<br>OR | SE or<br>95%CI | P | Beta SD | SE or<br>95% CI | P | GWAS<br>ref |
| Body fat % | 443000 | -0.051 | -0.0060 | 0.0015 | 0.00013 | 69791 | -0.0045 | 0.0063 | 0.48 | -0.0059 | 0.0015 | 0.00005 | [40] |
| BMI | 449325 | -0.017 | -0.0035 | 0.0021 | 0.1 | 223372 | -0.0096 | 0.0042 | 0.022 | -0.0047 | 0.0019 | 0.012 | [41] |
| Hip circ. | 450276 | -0.11 | -0.0122 | 0.0020 | 8x10<sup>-10</sup> | 134725 | -0.0210 | 0.0051 | 0.000049 | -0.0134 | 0.0019 | 7x10<sup>-13</sup> | [35] |
| Hip circ.* | 450276 | -0.09 | -0.0110 | 0.0023 | 0.0000019 | N/A |  |  |  |  |  |  |  |
| Waist circ. | 450323 | -0.05 | -0.0039 | 0.0018 | 0.022 | 143054 | -0.0120 | 0.0048 | 0.01 | -0.0049 | 0.0017 | 0.004 | [35] |
| WHRadjBMI | 449216 |  | 0.0101 | 0.0022 | 0.000004 | 129682 | 0.0120 | 0.0048 | 0.014 | 0.0104 | 0.002 | 2x10<sup>-7</sup> | [35] |
| WHR | 449216 |  | 0.0068 | 0.0022 | 0.0022 | 133877 | 0.0060 | 0.0048 | 0.21 | 0.0067 | 0.002 | 0.001 | [35] |
| Height | 450112 | -0.095 | -0.0103 | 0.0017 | 2x10<sup>-9</sup> | 239542 | -0.0100 | 0.0033 | 0.0021 | -0.010 | 0.0015 | 1x10<sup>-11</sup> | [42] |
| **Metabolic trait** |  |  |  |  |  |  |  |  |  |  |  |  |  |
| ACR | 437029 | 0.003 | 0.0121 | 0.0021 | 6x10<sup>-9</sup> | N/A |  |  |  |  |  |  |  |
| BP meds ** | 93036/354886 | N/A | 0.0034 | 0.0008 | 0.000039 | N/A |  |  |  |  |  |  |  |
| CAD ** | 37741/318892 | N/A | 0.99 | 0.97-1.00 | 0.043 | 60801/123504** | 1.000 | 0.98-1.02 | 0.99 |  |  |  | [43] |
| DBP | 449332 | 0.13 | 0.0092 | 0.0020 | 0.0000038 | 111783 | 0.111^ | 0.0460^ | 0.03 | 0.121^ | 0.0233^ | 2x10<sup>-7</sup> | [44] |
| SBP | 450075 | 0.29 | 0.0120 | 0.0018 | 2x10<sup>-10</sup> | 108620 | 0.157^ | 0.0734^ | 0.06 | 0.255^ | 0.0373^ | 9x10<sup>-12</sup> | [44] |
| SBP*** | 378880 | 0.34 | 0.0143 | 0.0024 | 2x10<sup>-9</sup> | N/A |  |  |  |  |  |  |  |
| SBPadj1**** | 288,247 | 0.38 | 0.0159 | 0.0027 | 3x10<sup>-9</sup> | N/A |  |  |  |  |  |  |  |
| SBPadj2**** | 284,360 | 0.37 | 0.0154 | 0.0027 | 1x10<sup>-8</sup> | N/A |  |  |  |  |  |  |  |
| Hypertens.** | 241691/206525 | N/A | 0.0051 | 0.0010 | 0.00000066 | N/A |  |  |  |  |  |  |  |
| T2D** | 14371/428017 | N/A | 1.00 | 0.98-1.03 | 0.79 | 26488/83964** | 1.01 | 0.97-1.04 | 0.72 | 1.006 | 0.99-1.03 | 0.51 | [45] |

WHR = waist hip ratio, adj=adjusted, ACR = albumin creatine ratio, SBP and DBP = systolic and diastolic blood pressure. Meds = medication. Circ. = circumference. CAD=Coronary artery disease, T2D= type 2 diabetes, Hypertens. = hypertension. N/A = data from variant not available in published GWAS. Beta raw refers to effect size in real units, cm, mmHg and kgm<sup>2</sup> as applicable. None of the two sample meta-analyses showed evidence of heterogeneity at p<0.05.
\*adjusted for height
\*\*number of cases and controls
\*\*\*in subset of data, excluding related individuals
\*\*\*\*adjusted1 Adjusted for self reported units alcohol per day. Adjusting for self reported frequency of alcohol consumption:(beta 0.014, se 0.002, p=2x10-9)
\*\*\*\*adjusted2 Adjusted for self reported units of alcohol per day, smoking and salt intake
^effect sizes from Wain et al in mmHg, p values GC corrected

### The minor allele at FGF21 rs838133 is associated with higher blood pressure and altered lipid and liver enzyme levels, but not type 2 diabetes or heart disease

The A allele at *FGF21,* associated with lower body fat, was also associated with higher blood pressure, hypertension and blood pressure medication use. The effect sizes were not reduced after correcting for self reported alcohol intake, smoking and salt intake (**Table 2**). There was no association with coronary artery disease or type 2 diabetes. We also noted an association with albumin creatine ratio (ACR) when treated as a continuous trait (**Table 2**). The largest effect was with systolic blood pressure, where each A allele raised systolic blood pressure by 0.29 mmHg.

Finally, we examined the association between the *FGF21* rs838133 allele and relevant glycaemic and liver and lipid markers that were not available in the UK Biobank, but were available in published Genome Wide Association Study (GWAS) data (**Table 3**). The allele associated with higher sugar intake and less fat in the lower body was also associated with higher LDL-cholesterol, triglyceride and Gamma-glutamyl transpeptidase (GGT) levels and lower Alkaline phosphatase (ALP) levels in existing GWAS data [16, 17]. The associations with GGT and ALP were consistent with an effect of higher alcohol intake and lower protein and fat intake, respectively. Details of alcohol and macronutrient intake are not available in these studies to confirm this as the cause of the liver function test associations.

**Table 3.** Associations between the minor allele at rs838133 and liver, lipid and glycaemic traits in published GWAS. Note that the variant is not present on the metabochip genotyping array, meaning sample sizes for these traits are appreciably smaller than those available in the UK Biobank. References for GWAS used are [16, 17, 24]

| Trait | N | BETA | SE | P | GWAS ref |
| --- | --- | --- | --- | --- | --- |
| ALP | 30846 | -0.0043 | 0.0013 | $2.7 \times 10^{-7}$ | [16] |
| ALT | 53682 | 0.0028 | 0.0018 | 0.30 | [16] |
| AST | 39020 | 0.0025 | 0.0022 | 0.25 | [16] |
| GGT | 55885 | 0.0084 | 0.0023 | $3.7 \times 10^{-6}$ | [16] |
| HDL-C | 92820 | 0.0007 | 0.0055 | 0.96 | [17] |
| LDL-C | 88433 | 0.027 | 0.0059 | $1.7 \times 10^{-5}$ | [17] |
| Triglycerides | 89485 | 0.0165 | 0.0052 | 0.002 | [17] |
| Fasting glucose | 51750 | 0.0021 | 0.0036 | 0.55 | [24] |
| Fasting insulin | 51750 | -0.0004 | 0.0036 | 0.92 | [24] |
| HbA1C | 51750 | -0.0059 | 0.0042 | 0.15 | [24] |
| HOMAB | 51750 | -0.0022 | 0.0037 | 0.55 | [24] |
| HOMAIR | 51750 | -0.0022 | 0.0046 | 0.63 | [24] |

ALP= alkaline phosphatase, GGT=Gamma-glutamyl transpeptidase, ALT=alanine aminotransferase, AST= aspartate aminotransferase, HDL-C = high density lipoprotein cholesterol, LDL-C = low density lipoprotein cholesterol, HOMA = homeostatic model assessment of B, beta cell and IR, insulin resistance.

### The phenotypes associated with common alleles in the KLB and PPARG genes contrast with those of the FGF21 allele

To assess further the potential mechanisms of the *FGF21* rs838133 A allele’s effects on nutrient intake, anthropometric and metabolic traits, we examined two other common human alleles in the UK Biobank. The *KLB* gene encodes B-Klotho, a coreceptor for FGF21[18] and it’s presence in neurons was recently reported to be essential for FGF21’s weight loss effects[19]. The *PPARG* gene encodes Peroxisome proliferator-activated receptor gamma, an adipocyte transcription factor that upregulates FGF21 in white adipose tissue of db/db mice[20, 21]. A common allele in *KLB* is associated with alcohol intake[22] and a common allele in *PPARG,* Pro12Ala, is associated with type 2 diabetes[23] and fasting insulin[24]. We reasoned that, if the *FGF21* allele’s effect acted through its interaction with B-Klotho or PPARG, we would see similar patterns of associations between these alleles and human traits as we did for the *FGF21* allele.

There were several similarities and differences between the three alleles. First, the *FGF21* and *KLB* alleles had different patterns of association with nutrient and alcohol intake. The *KLB* allele associated with higher self-reported alcohol consumption was associated with lower percentage carbohydrate intake, the opposite pattern to that of the *FGF21* allele. Second, the *FGF21* and *KLB* alleles had different patterns of association with body composition. The *KLB* allele associated with higher self-reported alcohol consumption tended to be associated with higher body fat percentage and markers of adiposity, the opposite pattern to that of the *FGF21* allele. Both *KLB* and *FGF21* alleles associated with higher self-reported alcohol consumption were associated with higher blood pressure, but the *FGF21* allele’s blood pressure effect was stronger than that of the *KLB* allele, despite a weaker effect on alcohol consumption (both differences p<0.05). The associations between the *FGF21* allele and body composition and blood pressure were more similar to those of the *PPARG* Pro12Ala allele. The body fat percentage lowering allele of each was associated with higher blood pressure, although the *PPARG* allele was associated with a more favourable WHR. All three alleles were associated with reduced height. Finally, we showed that the *KLB* variant’s pattern of associations is most similar to that of the variant in the *ADH* gene known to alter alcohol intake. Full details of the associations of all four variants are given in **supplementary table 3**.

### Phenome Wide Association Study shows suggestive associations of FGF21 allele with circadian rhythm and physical activity

FGF21 has multiple metabolic effects, possibly all linked to a role as a “global starvation signal”[25], and studies in mice have identified links to growth, bone metabolism[21], circadian rhythms[25], physical activity[25] and reproductive traits[26]. We therefore performed a “Phenome wide association study” (PheWAS), testing the association of the *FGF21* rs838133 variant with 105 traits in UK Biobank in the 451,000 individuals of European ancestry. We used a false discovery rate of 1% (p<0.0095) to highlight associations (full list of 105 associations in **supplementary table 4**). Notable associations, separate from the traits already mentioned, between the allele associated with higher sugar intake and other traits, included a more evening chronotype, lower activity (as measured by the international physical activity questionnaire (IPAQ)) and lower birth weight (all self reported). There was no association with bone mineral density as measured by a heel ultrasound, but the sugar intake allele was associated with a nominally higher risk of osteoporosis (**supplementary table 4**). There were no associations with the tested female reproductive traits. We further tested the associations with physical activity and sleep using a subset of 96,034 individuals who had worn accelerometer devices for 7 days and saw consistent associations, with nominal levels of statistical confidence (**supplementary table 5**).

### There is no evidence that the FGF21 allele alters FGF21 expression in liver

We looked up the association of the *FGF21* rs838133 variant and its association with gene expression in the GTEX database of human gene expression. There was no evidence this variant is associated with FGF21 gene expression in the liver based on 153 samples.

## DISCUSSION

The genotype-phenotype associations of the naturally occurring human *FGF21* variant rs838133 have potential implications for the development of FGF21 based therapies. Some of these therapies have reached early clinical trials and suggest that FGF21 mimetics can induce weight loss and reduce insulin resistance [7–9]. Whilst there is no direct evidence for the function of the rs838133 variant (or one in strong linkage disequilibrium), the A allele is very likely to represent a loss of FGF21 function, because it is very robustly associated with higher sugar and alcohol preference in people, a finding that is completely consistent with the genetic and pharmacological effects of FGF21 lowering in animal models, including non-human primates[1, 2]. The apparently paradoxical association between the *FGF21* A allele and higher circulating levels of FGF21 protein[12, 14], may be caused by a feedback process that results in greater FGF21 production as a response to higher carbohydrate intake.

The developers of therapies targeting FGF21 will be interested in the genotype-phenotype associations of the putative loss of function allele with a) higher percentage carbohydrate intake, b) higher alcohol intake, c) lower body fat percentage, d) altered body fat distribution, e) higher blood pressure, f) higher albumin to creatine ratio in the urine, g) higher LDL-cholesterol, h) higher triglycerides, but i) minimal if any effect on BMI and j) no evidence of an effect on type 2 diabetes or glycaemic traits. Of these associations, only that with higher percentage carbohydrate intake was convincingly shown before[13].

There are notable similarities and differences between the genotype-phenotype associations of the *FGF21* allele and the effects of FGF21 based interventions. The human genetic data are consistent with the reported beneficial effects of FGF21 mimetics on lipids and no, or limited, effect on diabetes and insulin sensitivity, although, for insulin sensitivity especially, null associations need to be interpreted with caution given the much smaller (~1/9^th^) sample sizes available for these traits compared to those in the UK Biobank. In contrast, the data suggest interventions that raise or deliver more FGF21 may not alter BMI, but may lower blood pressure and raise body fat in the lower body.

The multiple proposed effects of FGF21 led us to perform a “phenome wide association study”, of 105 traits in the UK Biobank. Some of the associations are consistent with those observed in studies of mice that suggest FGF21 is a “global starvation” signal. For example, the allele that increases sugar intake, is associated with lower levels of physical activity and altered chronotype, both features of mice with higher FGF21[25], but not with female reproductive traits or bone mineral density.

The effects of the *FGF21* allele are extremely small, at approximately 1/3^rd^ of one mmHg blood pressure and 1mm difference in hip circumference and height. However, the effect sizes of common genetic variants are not important when it comes to using them to provide insight into likely effects of much larger perturbations of a potential target. This point was recently illustrated by the very subtle but significant effects of common alleles in genes encoding lipid lowering proteins and type 2 diabetes [27–30].

There are two important caveats to using the human genetic associations to predict potential consequences of FGF21 based therapies. First, the associations with blood pressure and body fat distribution may be “off-target” – the consequences of the indirect effects on behaviour and eating habits, rather than “on-target” – the consequence of a direct effect of FGF21 altering blood pressure and body fat distribution. For example, the allele associated with higher sugar intake, is also associated with reward-based behaviours, including higher alcohol intake. These associations are consistent with studies showing signalling effects of FGF21 to the paraventricular hypothalamus and behavioural studies in mice and non-human primates[1, 2]. The *FGF21* allele’s association with blood pressure is not fully explained by altered alcohol intake, because the effects on blood pressure are larger than that of the *KLB* allele, and effects do not change when adjusting for self report alcohol intake. However, we cannot rule out other effects on reward based behaviours that could alter blood pressure. Second, the *FGF21* allele is very likely to influence human traits throughout life and therefore may have different effects compared to an acute, pharmacological based intervention.

The comparisons of the effects of the *FGF21* allele with those of the *KLB* and *PPARG* alleles provide further insight into potential mechanisms by which the allele might alter human traits. First, the data suggest that the *FGF21* allele’s effects on blood pressure and body composition are at least partly independent of any action through B-Klotho. Second, for body composition and blood pressure, the data are more consistent with the *FGF21* allele having a role similar to that of the *PPARG* allele, with the allele associated with lower body fat percentage, also being associated with adverse metabolic effects. Some studies have linked FGF21’s function in adipocytes to those of PPARG [21], a transcription factor critical for adipogenesis and mutations in which cause a form of lipodystrophy characterised by greatly reduced subcutaneous body fat, insulin resistance, high circulating triglyceride levels and higher blood pressure[31]. The human genetic associations suggest further studies are needed to investigate FGF21’s role in adipocyte differentiation and storage capacity. The *FGF21* rs838133 minor allele is not the first common allele to be associated with apparently paradoxical effects on fat mass and metabolic markers. Previous human genetic studies of alleles associated with insulin sensitivity have shown that most are also associated with an apparently paradoxical higher fat mass (and the insulin resistance allele is associated with lower fat mass)[32–37]. Like the *FGF21* allele, for some of these alleles the apparent paradox is explained by the higher fat mass being concentrated in the lower body, at least in women, (e.g. those in or near *FAM*, *LYPLAL1*, *GRB14*) [35].

There are two further limitations to our study that also mean caution is needed when translating the human genetic findings to implications for FGF21 therapies. First, despite being in the exon of *FGF21,* we cannot rule out the possibility that the variant associated with macronutrient intake operates through a nearby gene, and we note that the rs838133 allele is associated with the expression of a nearby gene *FUT2*, involved in vitamin B12 metabolism[38]. If this was the case it would mean our inferences about any pharmacological interventions targeting FGF21 are invalid. However the similarity between the human, mouse and non-human primate nutrient preference phenotypes suggests this is unlikely. Second, even if the variant affects *FGF21*, we do not know exactly how it alters sugar intake and it may do so in a way that disrupts the protein rather than simply changes its levels, for example through alternative splicing. However based on a bioinformatics analysis of predicted splicing motifs, the most likely causal variant, rs838133 is unlikely to alter *FGF21* splicing.

In summary, human genetic association data provides further insight into the potential multiple metabolic effects of FGF21 and has implications for the development of therapies targeting FGF21.

## Acknowledgements

This research has been conducted using the UK Biobank Resource. This work was carried out under UK Biobank project number 9072 and 9055. We thank Alexei Kharitonenkov Robert Andrews and Tim McDonald for helpful comments.

## Ethics UK Biobank

This study was conducted using the UK Biobank resource. Details of patient and public involvement in the UK Biobank are available online (www.ukbiobank.ac.uk/about-biobank-uk/ and https://www.ukbiobank.ac.uk/wp-content/uploads/2011/07/Summary-EGF-consultation.pdf?phpMyAdmin=trmKQlYdjjnQIgJ%2CfAzikMhEnx6). No patients were specifically involved in setting the research question or the outcome measures, nor were they involved in developing plans for recruitment, design, or implementation of this study. No patients were asked to advise on interpretation or writing up of results. There are no specific plans to disseminate the results of the research to study participants, but the UK Biobank disseminates key findings from projects on its website.

## Methods

### UK Biobank cohort

UK Biobank recruited over 500,000 individuals aged 37-73 years (99.5% were between 40 and 69 years) between 2006-2010 from across the UK. Participants provided a range of information via questionnaires and interviews (e.g. demographics, health status, lifestyle) and anthropometric measurements, blood pressure readings, blood, urine and saliva samples were taken for future analysis: this has been described in more detail elsewhere[39]. SNP genotypes were generated from the Affymetrix Axiom UK Biobank array (~450,000 individuals) and the UKBiLEVE array (~50,000 individuals). This dataset underwent extensive central quality control (http://biobank.ctsu.ox.ac.uk). We based our study on 451,099 individuals of white European descent as defined by Principal Components analysis (PCA). Briefly, principal components were generated in the 1000 Genomes Cohort using high-confidence SNPs to obtain their individual loadings. These loadings were then used to project all of the UK Biobank samples into the same principal component space and individuals were then clustered using principal components 1 to 4. We removed 7 participants who withdrew from the study, and 348 individuals whose self-reported sex did not match their genetic sex based on relative intensities of X and Y chromosome SNP probe intensity.

### Food frequency questionnaire (FFQ) and alcohol intake in UK Biobank participants

The food frequency questionnaire (FFQ) was added towards the end of the recruitment phase and participants completed whilst at the recruitment centre. Participants were then sent 4 FFQs and asked to complete online. The questionnaire focussed on the consumption of approximately 200 commonly consumed food and drinks (http://biobank.ctsu.ox.ac.uk/crystal/refer.cgi?id=118240). For each participant completing the food frequency questionnaire nutrient intakes were estimated by multiplying the quantity consumed by the nutrient composition of the food or beverage, as taken from the UK food composition database McCance and Widdowson’s The Composition of Foods and its supplements (FSA, 2002). 211,051 participants completed at least one FFQ. Participants were asked if this was a standard diet day for them and we excluded the 18,054 participants who reported not following a standard diet. Averages were then calculated for participants with up to five normal questionnaires for 192,997 individuals.

We derived alcohol units consumed per day for individuals in the UK Biobank. For individuals reporting drinking alcohol at least once a week a units per week variable was calculated and for individuals reporting less frequent drinking a units per month variable was calculated. A 125ml glass of wine (red, white or sparkling) was considered to be 1.5 units, a pint of beer or cider was considered to be 2.8 units, other alcoholic drinks (e.g. alcopops) was considered to be 1.5 units and a measure of spirit was considered to be 1 unit.

### Coffee and smoking and salt intake in UK Biobank participants

All participants in the UK Biobank were asked about their smoking status, with individuals defined as never, former or current smokers. All participants in the UK Biobank were asked about adding salt to food. Participants were asked: “Do you add salt to your food? (Do not include salt used in cooking)”, with the options “Never/rarely”, “Sometimes”, “Usually” or “Always”. In the general questionnaire participants were asked "How many cups of coffee do you drink each DAY? (Include decaffeinated coffee)". Participants were also asked about whether they drank caffeinated or decaffeinated coffee. From this we derived a number of cups of coffee per day and a number of cups of caffeinated coffee per day.

### Disease and related anthropometric and metabolic traits in UK Biobank participants

#### Measures of adiposity

We used Bio-impedance measures of body fat percentage measured by the Tanita BC418MA body composition analyser.

#### Measures of disease and disease related traits

We defined type 2 diabetes, hypertension, blood pressure and heart disease using baseline data and following similar definitions to those used in previous genome wide association studies. For coronary artery disease we additionally included cases from hospital episodes statistics available at the time (through 31^st^ March 2016 release, ICD10 codes I21*, I22*, I23*, I24* and I25*). We defined type 2 diabetes cases if 3 criteria were present: i) reports of either type 2 diabetes or generic diabetes at the interview, ii) at least one year gap from diagnosis without requiring insulin iii) reported age at diagnosis over the age of 35 years to limit the numbers of individuals with slow-progressing autoimmune diabetes or monogenic forms. Individuals not reporting an age of diagnosis were excluded. We also excluded individuals diagnosed with diabetes within the year prior to the baseline study visit as we were unable to determine whether they were using insulin within the first year. Controls were individuals not fulfilling these criteria.

We defined hypertensive cases as individuals with systolic blood pressure of >140 mmHg, or a diastolic blood pressure of >90 mmHg, or the report of blood pressure medication usage. Controls were individuals not fulfilling these criteria. For the analysis of systolic and diastolic blood pressure, we corrected blood pressure measures in people on antihypertensive drugs by adding 15 mmHg to systolic and 10 mmHg to diastolic blood pressure, in keeping with the approach taken by genome wide association studies.

We defined heart disease cases if individuals reported angina and/or a heart attack at the interview stage. We defined Controls as individuals without these conditions.

### Traits derived from accelerometers

The UK Biobank has collected accelerometer data in 103,711 participants, who wore the devices on their wrist for a continuous period of a week. We used a well-validated and freely available R package called GGIR (v1.5-12) to process these files, made available to researchers, in order to extract measures of physical activity and sleep. For this study, we used three derived measures of activity and one measure of sleep timing (L5 time). L5 time represents the midpoint time of the least active five hours of the day, as defined by the minimum point of a moving average of activity levels. Our L5 time variable represents an individual’s average across all days recorded and units are reported in number of hours after previous midday. In addition, we also used measures of physical activity from the UK Biobank to define the proportion of activity classified as 1) sedentary or asleep (<40 milligravities (mg)), 2) the proportion of activity at least non-sedentary (>40mg), and 3) the proportion of activity over a threshold representing moderate-to-vigorous activity (>100mg).

### Traits in published GWA studies

We looked up the association of the variant rs838133 in existing relevant genome wide association studies, as detailed in the tables. The variant was not present on the metabochip.

### Statistical analysis

All genotype-phenotype association data were generated starting from 451,099 individuals defined as European ancestry and using BOLT-LMM v1.2, that uses an LD score regression approach to account for structure caused by relatedness (close and distant) [REF]. All association testing was based on an additive, per allele, model and adjusted for SNP chip type (UKB Axiom or UK BiLEVE), test centre, sex and age (or year of birth for age at menarche). Accelerometry based phenotypes were additionally adjusted for season and age at wear time. We tested approximately 100 traits and so highlight main associations reaching p<0.0005, but, given several metabolic and anthropometric traits reach genome wide significance, and the known role of the FGF21 variant, we mention other traits reaching nominal significance and used a false discovery rate of 1% to highlight associations in the PheWAS. For continuous traits, we inverse normalised phenotypes to account for any skewed distributions,

**Figure 1.**
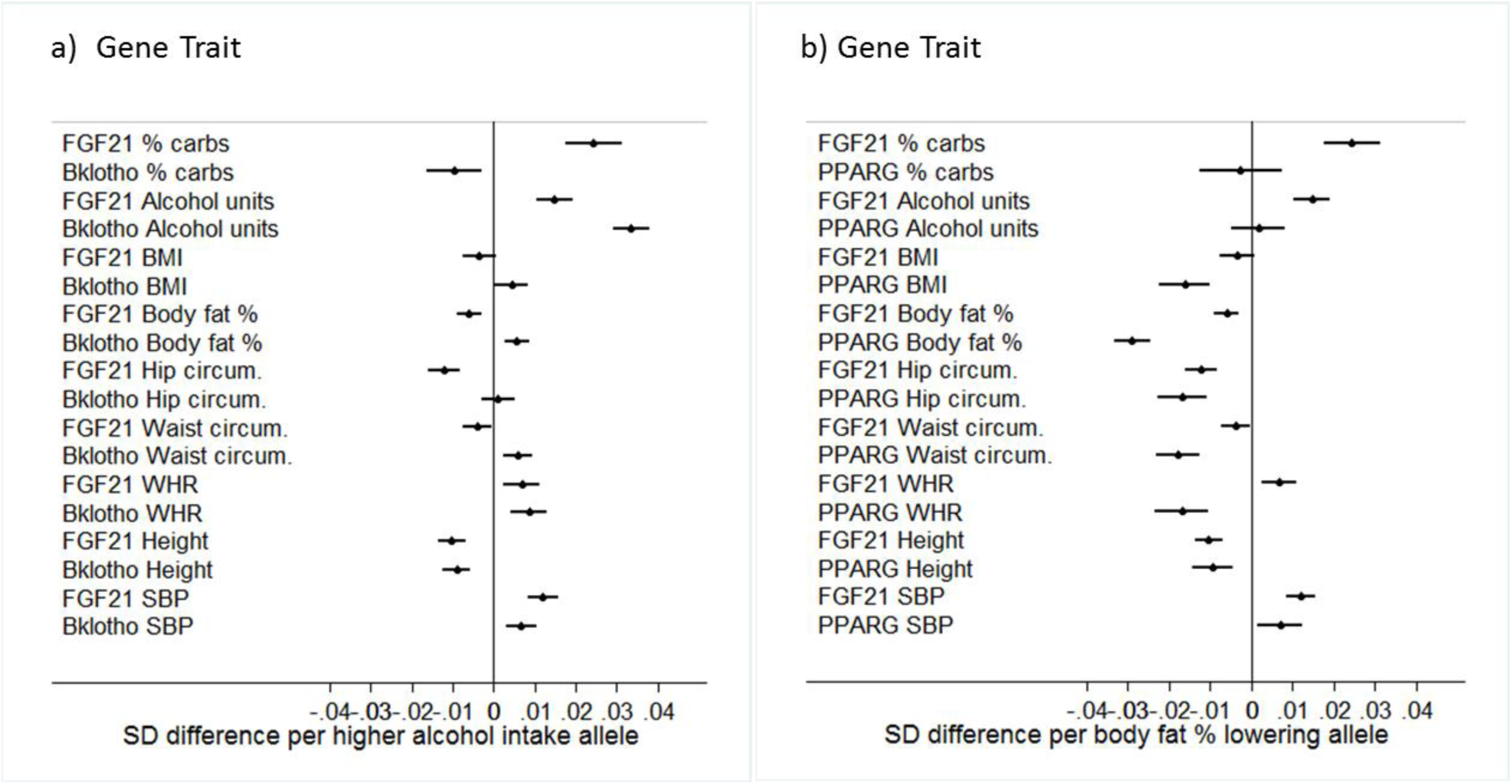
Genotype phenotype associations of the *FGF21* rs838133 A allele compared to a) the *KLB* (Bklotho) rs28712821 A allele and b) the *PPARG* rs1801282 (Pro12Ala) C allele in the UK Biobank. SBP=systolic blood pressure, DBP=diastolic blood pressure, WHR=waist hip ratio. Effects are orientated to the higher alcohol intake allele for the comparision with KLB, and the body fat % lowering allele for PPARG.

